# Real-time analysis and visualization of pathogen sequence data

**DOI:** 10.1101/286187

**Authors:** Richard A. Neher, Trevor Bedford

## Abstract

The rapid development of sequencing technologies has to led to an explosion of pathogen sequence data that are increasingly collected as part of routine surveillance or clinical diagnostics. In public health, sequence data is used to reconstruct the evolution of pathogens, anticipate future spread, and target interventions. In clinical settings whole genome sequences identify pathogens at the strain level, can be used to predict phenotypes such as drug resistance and virulence, and inform treatment by linking to closely related cases. While sequencing has become cheaper, the analysis of sequence data has become an important bottleneck. Deriving interpretable and actionable results for a large variety of pathogens – each with their own complexities – from continuously updated data is a daunting task and requires flexible bioinformatics workflows and dissemination platforms. Here, we review recent developments in real-time analysis of pathogen sequence data with a particular focus on visualization and integration of sequence and phenotypic data.

---

As pathogens replicate and spread, their genomes accumulate mutations. These changes can now be detected via cheap and rapid whole genome sequencing on unprecedented scale. Such sequence data are increasingly used to track the spread of pathogens and predict their phenotypic properties. Both applications have great potential to inform public health and treatment decisions if sequencing data can be obtained and analyzed rapidly. Historically, however, sequencing and analysis has lagged months-to-years behind sample collection. The results from these studies have taught us much about pathogen molecular evolution, genotype-phenotype maps, and epidemic spread, but have come almost always too late to inform public health interventions or treatment decisions.

The rapid development of sequencing technologies has made routine sequencing of viral and bacterial genomes possible and tens of thousands of whole genome sequences (WGS) are deposited in databases every year (see Fig. 1). Many more, regrettably, are sequenced and not shared. There are currently two major directions in which high-throughput sequencing technologies are used in public health and diagnostics: (i) to track outbreaks and epidemics to inform public health response, and (ii) to characterize individual infections to tailor treatment decisions.

**FIG. 1.**
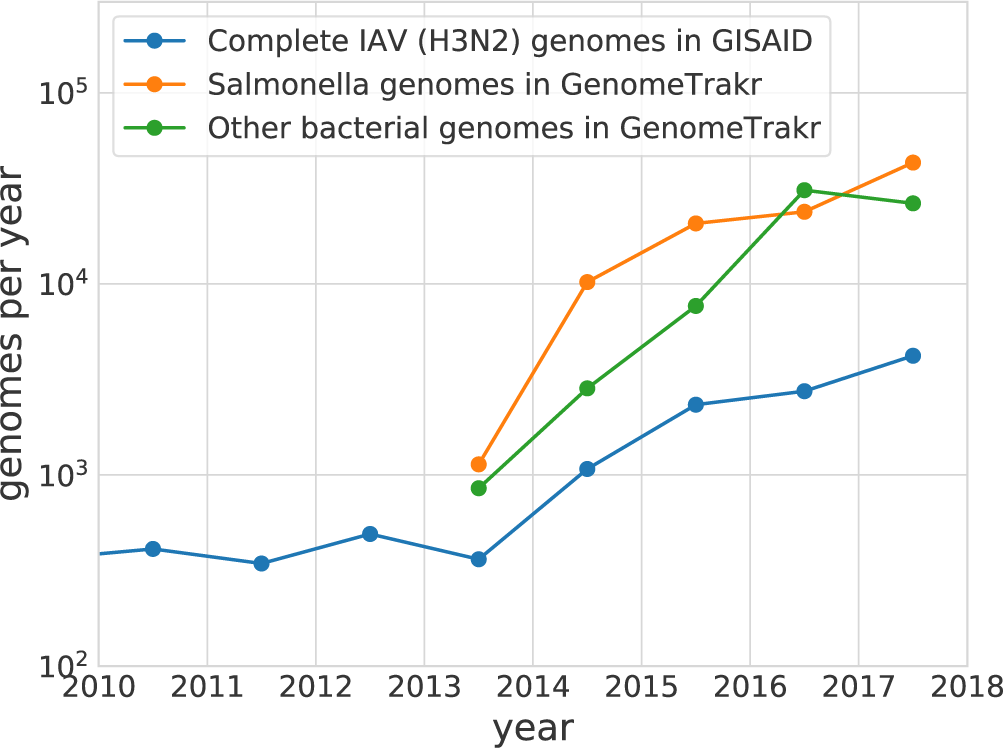
The number of complete pathogen genomes has increased dramatically over the last few years. More than 4000 complete influenza A (IAV) subtype H3N2 virus genomes have been deposited in GISAID in 2017. The GenomeTrakr network sequenced in excess of 40,000 Salmonella genomes and 25,000 other bacterial genomes (mostly *Listeria*, *E. coli/Shigella*, and *Campylobacter*) in 2017^11^.

## Sequencing in public health

The utility of rapid sequencing and phylogenetic analysis of pathogens is perhaps most evident for influenza viruses and food-borne diseases. Due to rapid evolution of its viral surface proteins, the antigenic properties of the circulating influenza viruses change every few years and the seasonal influenza vaccine needs frequent updating^1^. The WHO Global Influenza Surveillance and Response System (GISRS) sequences hundreds of viruses every month and many of these sequences are submitted to the GISAID database (gisaid.org) within 4 weeks of sample collection. Phylogenetic analysis of these data provide an accurate and up-to-date summary of the spread and abundance of different viral variants that is crucial input to the biannual consultations on seasonal influenza vaccine composition.

Such rapid turn-around and data sharing is considerably harder to achieve in an outbreak setting in resource limited conditions. Nonetheless, Quick et al.^2^ achieved even shorter turn-around during the tail end of the 2014–2015 West African Ebola outbreak. Similarly, Dyrdak et al.^3^ analyzed an enterovirus outbreak in Sweden and continuously updated the manuscript until publication with sequences sampled within days of publication included in the analysis.

Molecular epidemiology techniques can reconstruct the temporal and spatial spread of an outbreak. In this case, the accumulation of mutations alongside a molecular clock estimate can be used to date the origin of an outbreak. Similarly, by linking samples that originate from different geographic locations, phylogeographic methods can reconstruct geographic spread and differentiate distinct introductions. The resolution of these inferences critically depends on the rate at which mutations accumulate in the sequenced locus, which increases with the per site evolutionary rate and the length of the locus.

RNA viruses accumulate changes in their genome with a typical rate of 0.0005 to 0.005 changes per site and year^4^. Rate estimates vary from virus to virus and depend on the time scale of observation or whether measured within or between hosts. Ebola virus and Zika virus, for example, evolve at a rate of *µ* ≈ 0.001 per site per year. The expected time interval without a substitution along a transmission chain is 1/(*µL*), which corresponds to approximately 5 weeks for Zika virus (*L* ≈ 10kb) and 3 weeks for Ebola virus (*L* ≈ 19kb). Hence evolution and spread of such RNA viruses can be resolved on the scale of a month. While this temporal resolution is typically insufficient to resolve individual transmissions, it is high compared to the duration of outbreaks. Rapid sequencing and analysis therefore has the potential to inform intervention efforts as outbreaks are unfolding. In particular, they rule out direct transmission and differentiate different introductions or zoonosis.

Phylodynamic and phylogeographic methods are best established for viral pathogens with high evolutionary rates and small genomes for which large scale sequencing has been possible for years. The evolutionary rates of bacteria are many orders of magnitudes lower than those of RNA viruses. However, bacteria also have about 100 to 1000-fold larger genomes and it is now possible to sequence entire bacterial genomes at low cost. Substitution rate estimates in bacteria come with substantial uncertainty but they tend to be on the order of one substitution per megabase per year (with about one to two orders of magnitude of variation between species^5^). With a typical genome size of 5 megabases, this translates into 5–10 substitutions per genome and year — similar to many RNA viruses. The substitution rate in the core genome of MRSA, for example, was estimated to be 1.3 × 10^*−6*^ per site and year^6^. The core genome of *Listeria monocytogenes* evolves more slowly at about one substitution every 2.5y^7^. Hence real-time phylogenetics for bacterial outbreak tracking is possible in much the same way as for RNA viruses. Analysis of bacterial genomes, however, is vastly more complicated than that of RNA viruses with short genomes. Bacteria frequently exchange genetic material via horizontal transfer, take up genes from the environments and rearrange their genome. Recombination can blur phylogenetic signal and recombinant sequence is often difficult to remove. Furthermore, strong selection within hosts, for example through drug therapy, can accelerate evolution by an order of magnitude^8^. If not properly accounted for, these processes can blur any temporal signal and obscure links between closely related isolates.

Even with whole genomes, phylogenetic resolution typically is insufficient to make the case for a direct transmission, but transmission can be confidently ruled out for divergent sequences, seemingly unrelated cases can be grouped into outbreaks (e.g. an outbreak of drug resistant MtB among migrants arriving in multiple European countries^9^), predominant routes of transmission and likely sources in the environment or animal reservoirs can be identified. Genome-Trackr and PulseNet, for example, are a large federated efforts to sequence tens of thousands of genomes from food-borne outbreaks and clinical samples^10;11^. All sequence data from these projects are publicly available on NCBI with little delay and are analyzed in real-time to track outbreaks. The recently released Pathogen Detection system by NCBI (www.ncbi.nlm.nih.gov/pathogens/) provides convenient access to the sequence and metadata generated by these projects as well as phylogenetic analysis.

These examples illustrate the potential and feasibility of obtaining actionable information from pathogen sequence data for both viral and bacterial infections. However, with rapidly increasing data volumes, efficient processing pipelines and tools that help with interpretation – e.g. visualizations – increasingly become the bottleneck.

## Sequencing in diagnostics and therapy

For some pathogens like Zika virus, sequencing the genome has no implications for treatment. In the case of HIV, however, drug resistance profiles derived from sequence data have directly informed treatment for years^12^. As the genetic basis of drug resistance phenotypes are better understood, rapid whole genome sequencing will increasingly be used to diagnose and phenotype pathogens directly from the clinical specimen. Such culture-free methods are particularly important for tuberculosis, in which culture based susceptibility testing takes many weeks. Votintseva et al.^13^ have recently shown that high-throughput sequencing directly from respiratory samples can provide drug resistance profiles of *M. tuberculosis* within a day.

Sequencing for diagnostic purposes or for public health surveillance have different aims and requirements, but can complement each other. Public health response typically requires recent data with an emphasis on dynamics. Surveillance data provides context for the individual case in the clinics requires a stable database with validated content to make reliable predictions on drug susceptibility, phylogenetic context, and protective measures. Clinical sequencing data, however, should be fed into surveillance databases immediately whenever ethically possible. Only with rapid and open sharing of sequencing data can the full potential of molecular epidemiology be realized^11^.

The challenges involved in sample collection, processing, sequencing and data sharing have been discussed at length elsewhere^14^. Here, we focus on software developments that facilitate the implementation of real-time analysis with an emphasis on web-based visualization, as a full review of general tools for genomic analysis and visualization is not easily encompassed.

## RAPID AND INTERPRETABLE ANALYSIS OF GENOMIC DATA

A typical molecular epidemiological analysis aims to identify transmission clusters, date the introduction of the pathogen, detail geographic spread, and in some cases identify potential phenotypic change of a pathogen from sequence data. The rapidly increasing numbers of sequenced genomes make comprehensive analysis computationally challenging. While 1000s of viral genomes can be aligned within minutes (e.g. by MAFFT) and the reconstruction of a basic phylogenetic tree typically takes less than one hour (e.g. using IQ-TREE, RAxML or FastTree), the most popular tool for phylodynamic inference (BEAST)^15^ will often take weeks to finish.

To overcome these hurdles, several tools that use simpler heuristics have been developed to infer time-stamped phylogenies^16;17;18^. Rather than sampling a large number of tree topologies, these tools use the topology of an input tree with little or no modification. Dating of ancestral events tends to be of comparable accuracy to BEAST^16;17;18^. However, these tools do not integrate uncertainty of tree reconstruction and provide limited flexibility to infer demographic models. Furthermore, the heuristics used by these program are based on assumptions (for example that sequences are closely related) and they are not expected to be accurate in all scenarios. The computational cost of Bayesian phylodynamics could be mitigated if methods for continuous updating and augmenting of the Markov chain with additional data were developed. For the time being, however, efficient heuristics and sensible approximations deliver sufficiently accurate and reliable results when near real-time analysis is required.

### Viral genomes: Nextflu and Nextstrain

The number of influenza viruses that are sequenced and phenotyped per month has increased sharply to a point that a comprehensive and timely manual analysis and annotation of the results is no longer feasible. In 2014, we developed an automated phylodynamic analysis pipeline that operates on an up-to-date database of sequences and serological information. The results of this pipeline are available were made available at nextflu.org and included a phylogeny, branch-specific mutations, frequency trajectories of mutations and variants, and a model of antigenic evolution.

Nextflu is now integrated in the more general platform Nextstrain, that provides an online platform for outbreak investigations of diverse viruses and is available at nextstrain.org^**?**^. Nextstrain uses TreeTime^18^ to infer timescaled phylogenies and conduct ancestral sequence inference. In addition, Nextstrain uses the discrete ancestral character inference of TreeTime to infer the likely geographic state of ancestral nodes. Since this approach applies “mutation” models to “migration”, it is often called a “mugration model”. A phylodynamic/phylogeographic analysis of 1000 sequences of length 10kb takes on the order of an hour on a standard laptop computer.

### Bacterial WGS data

Bacterial WGS data typically comes in the form of millions of short reads that can either be assembled into contigs, mapped against reference sequences, or classified based on kmer distributions. A large number of tools have been developed for rapid species identification, typing, and variant calling. WGSA by the, for example, allows the user to upload an assembly and WGSA will detect the species and infer the multi-locus sequence type within a few seconds. In addition, WGSA predicts antibiotic resistance profiles for a number of species. WGSA was developed by the Center for Genomic Pathogen Surveillance and is available at wgsa.net.

Bacterial genomes are very dynamic and frequently gain or lose genes. Even closely related bacteria can differ in the presence or absence of dozens of genes. To track transmission and detect clusters, genomes are typically compared at a set of *core genes* present in all bacteria of a species. Genes present in only a fraction of individuals are referred to as *accessory genes*.

Clinically important genes such antibiotic resistance determinants or virulence factors are often not part of the core genome and are horizontally transferred between strains and species. Collections of bacterial genomes are therefore analyzed using pan-genome tools that aim to cluster all genes in the collection of genomes into orthologous groups. Early methods for pan-genome analysis scaled poorly with the number of genomes that are analyzed since every gene in every genome needed to be compared to every other gene. The first tool capable of analyzing 100s of bacterial genomes was Roary^20^. Roary is designed to work with very similar genes (as is common in outbreak scenarios) and accelerates inference of orthologous gene clusters by pre-clustering genomes. A more recent pan-genome analysis pipeline capable of large scale analysis is panX^21^ that speeds up clustering by hierarchically building up the complete pan-genome from sub-pangenomes inferred from smaller batches of genomes. PanX is coupled to a web-based visualization platform discussed below.

While the pan-genome tools cluster annotated genes in the collection of genomes, they are of little help to assess the origin and distribution of a particular sequence. Traditional tools for homology search in NCBI only index assembled sequence, but today the majority of sequence data are stored in short read archives rather than in Genbank. Bradley et al.^22^ developed a method to search the entire collection of microbial sequence data including metagenomic samples from a wide variety of environment. The ability to search this vast treasure trove of data will likely be transformative in assessing spread and prevalence of novel resistance determinants. The recently discovered mobile colistin resistance gene *mcr-1*, for example, was found in more than 100 datasets where it wasn’t previously described^22^.

### Outlook

Most current analysis pipelines require rerunning the entire analysis even when only a single sequence is added. While this strategy is still feasible today, this will likely become unsustainable in the future. Applications that support cheap updating of datasets and on-line addition of user data will likely replace current versions.

## VISUALIZATION AND INTERPRETATION

With increasing dataset sizes, interpretation and exploration of data become increasingly challenging. Phylogenetic trees can be visualized as familiar planar graphs, but the tree alone only shows genetic similarity between isolates and becomes quickly unintelligible as the number of sequences increases. To make pathogen sequence data truly useful, it needs to be integrated with other types of information, ideally in an interactive way. A suitable platform to do so is the web-browser and several powerful web applications have emerged over the last few years. In addition, browser-based visualizations are naturally disseminated online.

### Microreact

Microreact is a web application based on React (a JavaScript framework for interactive applications), D3.js (a JavaScript library for producing dynamic, interactive data visualizations), Phylocanvas (a JavaScript flexible tree viewer), and Leaflet (a JavaScript mapping toolkit)^23^. Microreact allows exploration of a phylogenetic tree, the geographic locations, and a time line of the samples. It is available at microreact.org. Custom data sets can be loaded into the application in the form of a Newick tree and a tabular file containing information such as geographic location or sampling data.

### Nextstrain

Nextstrain was developed as a more generic and flexible version of Nextflu^19^ which is available at nextstrain.org. Similar to Microreact, Nextstrain uses React, D3.js, and Leaflet, but uses a custom made tree viewer that has flexible zooming and annotation options. The tree can be decorated with any discrete or continuous attribute, both on tips of the tree and inferred values for internal nodes (for example geographic location). Nextstrain maps individual mutations to branches in the tree and thereby allows to associate mutations with phenotypes or geographic distributions. The map in Nextstrain shows putative transmission events and a panel indicates genetic diversity across the genome (Fig. 2).

**FIG. 2.**
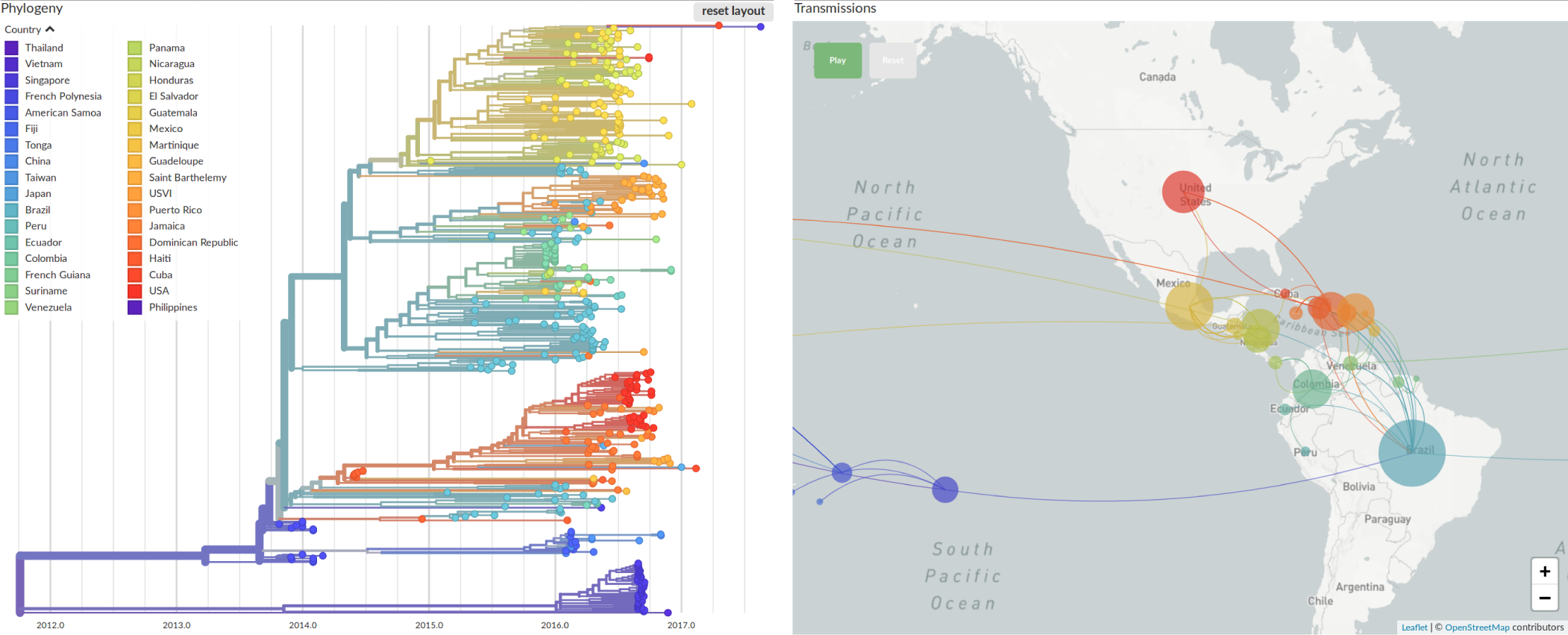
Phylogeographic analysis of Zika virus sequences on nextstrain.org?. Whole genomes sequences sampled between 2013 and 2017 were processed using the Nextstrain pipeline. Nextstrain reconstructs likely time and place of each internal node of the tree and from this assignments infers possible transmission patterns that are displayed on a map. Molecular analysis of this sort reveals for example multiple introductions of Zika virus into Florida originating most likely from viruses circulating in the Caribbean in 2015-2016.

The analyses by Nextstrain and Nextflu critically depend on timely and open sharing of sequence information that many laboratories around the globe contribute. To incentivize early pre-publication sharing of data, platforms like Nextstrain need to explicitly acknowledge the individual contributions. Ideally, such platforms should provide added value to authors, such as for example deep links that show data by a particular group in the context of the outbreak.

### Phandango

Phandango is an interactive viewer for bacterial whole genome sequencing data^24^ and combines a phylogenetic tree with metadata columns and gene presence-absence maps or recombination events. Phandango is available at phandango.net and can ingest the output of a number of analysis tools commonly used for the analysis of bacterial WGS data such Gubbins, Roary and BRAT.

### panX

PanX is a pan-genome analysis pipeline that is coupled to a web-browser based visualization^21^. Similar to Phandango, it displays a core genome SNP phylogeny but is otherwise more centered on genetic variation in individual genes. Pangenomes of about 100 bacterial species based on curated reference genomes are available at pangenome.de. The tree and alignment of each gene in the pan-genome can be accessed rapidly by a searching a table of gene names and annotations. PanX then displays gene and species tree side by side and maps gene gain and loss events to branches in the core genome tree and mutations to branches in the gene tree. Trees can be colored by arbitrary attributes such as resistance phenotypes and associations between genetic variation and these phenotypes can be explored.

### Other tools

SpreaD3 allows of visualization of phylogeographic reconstructions from models implemented in the software package BEAST^25^. PhyloGeoTool is a web-application to interactively navigate large phylogenies and to explore associated clinical and epidemiological data^26^. TreeLink displays phylogenetic trees alongside metadata in an interactive web application^27^.

## CHALLENGES IN DATA INTEGRATION AND VISUALIZATION

With rapidly increasing volumes of sequence data, decisions as to how the data are filtered and what analysis are shown become increasingly important. Epidemiological investigations of a novel outbreak typically seek to identify the sources, track the spread, and detect transmission chains. In this case, a generic combination of map, tree, and time line will often be an appropriate and sufficient visualization. Nextstrain and Microreact both follow this paradigm.

However, when analyzing established pathogens that continuously adapt to treatments, vaccines, or pre-existing immunity, more specific applications will be necessary since case data, phenotype data, and clinical parameters differ wildly by pathogen. Such data will generally have a common core such as sample date and location, but other parameters such as drug resistance phenotypes, disease severity, host age, risk group, serology, etc., are pathogen specific. While these data are often stored in non-standardized formats and ethical and technical reasons can impede data sharing, these data are often at least as important for interpretation of the epidemiological dynamics as phylodynamic inference from sequence data. The value of either data is greatly increased by seamless integration, but the idiosyncrasies require flexible analysis and visualization frameworks that can be tailored to specific pathogens.

One such example is the serological characterization of influenza viruses via hemagglutination inhibition (HI) titers using antisera raised in ferrets. Such titers are routinely measured as part of GISRS to monitor the antigenic evolution of influenza viruses are a good example how phenotype information can be interactively integrated with phylogeny and molecular evolution. HI titers are reported in large tables and have been traditionally visualized using multidimensional scaling without any reference to the phylogeny. In Bedford et al.^28^ and Neher et al.^29^, we developed methods to integrate the molecular and antigenic evolution of influenza virus. This integration allows association of genotypic changes with antigenic evolution and suggests an intuitive and interactive visualization of HI titer data on the phylogeny. A screenshot of this integration is shown in Fig. 3. Due to data sharing restrictions, most HI titer data are not openly available, but historical data by McCauley and colleagues are visualized along with the molecular evolution at hi.nextflu.org.

**FIG. 3.**
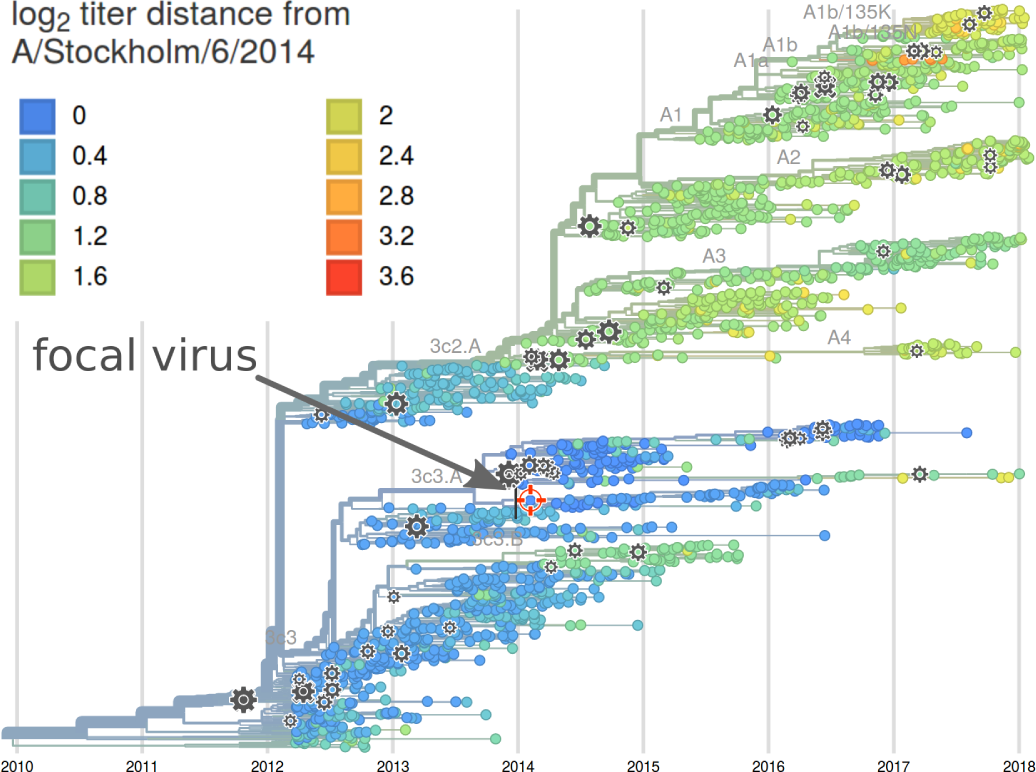
Integration of HI titer data with molecular evolution influenza virus. Each year, influenza laboratories determine thousands of HI titers of test viruses relative to sera raised against several reference viruses (indicated by gray cogs). These data can be integrated with the molecular evolution of the virus and visualized on the phylogeny (here showing inferred titers using a model). The reference virus with respect to which titers are displayed can be chosen by clicking on the corresponding symbol in the tree^29^. The visualization exposes both raw data (via tool tips for each virus) as well as a model inference that integrates many individual measurements (hi.nextflu.org).

In addition to phenotype integration, it is crucial to choose the right level of detail for a specific application. This is particularly true for bacteria where the relevant information might be the core genome phylogeny, the presence/absence of particular genes or plasmids, or individual mutations in specific genes. If the analysis tool and the visualization does not provide a fine grained analysis at the relevant level, the most important patterns might stay hidden. On the other hand, sifting through every gene or mutation is prohibitive. The primary aim should be to highlight the most important and robust patterns upfront and provide flexible methods to filter and rank variants (e.g. by recent rise in frequency, association with host, resistance or risk group, etc). The user should then have the possibility to expose detail on demand when a deeper exploration is required.

Similarly, parameter inferences and model abstractions are very useful to get a concise summary of the data, but should be complemented by the ability of interrogate the raw data (e.g. an estimate of the evolutionary rate should be accompanied by a scatter plot of root-to-tip divergence and sampling time). This is particularly important in outbreak scenarios when methods are applied to an emerging pathogen in a developing situation.

For clinical applications, the presentation of the results of an analysis should be focused on the sample in question and only provide reliable and actionable information, while suggestive and correlative results tend to be a distraction^30^.

## CONCLUSIONS

High-throughput and rapid sequencing is revolutionizing infectious disease diagnostics and epidemiology. Sequence data can be used to unambiguously identify pathogens, to link related cases, to reconstruct the spread of an outbreak, and will soon allow detailed prediction of a pathogen’s phenotype.

The Global Influenza Surveillance and Response System (GISRS) is a good example of a near real-time surveillance system. Hundreds of viruses are sequenced and phenotyped every month and the sequence data is shared in a timely manner. A global comprehensive analysis of these data, updated about once a week, is available at nextflu.org. These analysis directly inform the influenza vaccine strain selection process^1^. Several public health agencies have adopted WGS as their primary tool for outbreak investigation and many centers share these data openly with commendable timeliness. The GenomeTrakr and PulseNet networks, for example, now sequence and openly release about 5000 bacterial genomes per month^10;11^. These data are accessible on NCBI through the recently released Pathogen Detection system at www.ncbi.nlm.nih.gov/pathogens with analysis results available via FTP.

These two examples clearly show that near real-time genomic surveillance is possible and valuable and all the individual components to implement such surveillance are in place. However, to realize this potential for many more pathogens, sample collection and sequencing has to be streamlined, data need to be shared along with the relevant metadata, and specific analysis methods and visualizations need to be implemented and maintained.

## ACKNOWLEDGMENTS

We are indebted to scientists around the globe for timely and open sharing of sequence data, without which real-time visualization would not be worthwhile. We are grateful to James Hadfield for critical reading of the manuscript and Greg Armstrong for valuable feedback and insights into WGS surveillance efforts of bacteria.

